# Inducible lncRNA transgenic mice reveal continual role of HOTAIR in promoting breast cancer metastasis

**DOI:** 10.1101/2022.04.21.488980

**Authors:** Qing Ma, Liuyi Yang, Karen Tolentino, Yang Zhao, Ulrike M Lizenburger, Quanming Shi, Lin Zhu, Miao-Chih Tsai, Jun-An Chen, Ian Lai, Hong Zeng, Lingjie Li, Howard Y. Chang

## Abstract

HOTAIR is a 2.2 kb long noncoding RNA (lncRNA) whose dysregulation has been linked to oncogenesis, defects in pattern formation during early development, and irregularities during the process of epithelial-to-mesenchymal transition (EMT). However, the oncogenic transformation determined by HOTAIR in vivo and its impact on chromatin dynamics are incompletely understood. Here we generate a transgenic mouse model with doxycycline-inducible expression of human HOTAIR in the context of the MMTV-PyMT breast cancer-prone background to systematically interrogate the cellular mechanisms by which human HOTAIR lncRNA acts to promote breast cancer progression. We show that sustained high levels of HOTAIR over time increased breast metastatic capacity and invasiveness in breast cancer cells, promoting migration and subsequent metastasis to the lung. Subsequent withdrawal of HOTAIR overexpression reverted the metastatic phenotype, indicating oncogenic lncRNA addiction. Furthermore, HOTAIR overexpression altered both the cellular transcriptome and chromatin accessibility landscape of multiple metastasis-associated genes and promoted epithelial to mesenchymal transition. These alterations are abrogated within several cell cycles after HOTAIR expression is reverted to basal levels, indicating an erasable lncRNA-associated epigenetic memory. These results suggest that a continual role for HOTAIR in programming a metastatic gene regulatory program. Targeting HOTAIR lncRNA may potentially serve as a therapeutic strategy to ameliorate breast cancer progression.

## Introduction

Long noncoding RNAs (lncRNAs) are extensively transcribed from mammalian genomes, and may actively participate in diverse biological processes through means beyond protein production (Rinn and Chang, 2012). Tens of thousands of lncRNAs have been identified by high-throughput RNA sequencing, but only a small percentage of these species have been functionally characterized (Cabili et al., 2011; Consortium et al., 2014; Djebali et al., 2012; Iyer et al., 2015; Liu et al., 2021). Once thought to be byproducts of mRNA splicing with no known biological activity within the cell, a handful of lncRNAs have been demonstrated to act in regulating gene expression as well the deposition of epigenetic modifications (Kashi et al., 2016). Several lncRNAs serve important roles in gene regulation, including modulating gene transcription; controlling nuclear architecture; mRNA stability, translation and deposition of post-translational modifications (Yao et al., 2019). LncRNAs perform these functions through a variety of mechanisms, such as by acting as molecular scaffolds to bring multiple proteins into proximity in three-dimensional space; as ‘guides’ like Xist and HOTAIR which recruit chromatin-modifying enzymes to the genome; and as bridges which link promoters to distal enhancers to alter gene expression levels (Gupta et al., 2010; Lee, 2009; Rinn et al., 2007; Tsai et al., 2010). Some lncRNAs, such as LUNAR1 and CCAT1, facilitate or inhibit long-range chromatin interactions (Ma et al., 2015; Trimarchi et al., 2014; Xiang et al., 2014). The disruption of lncRNAs has been linked to the pathogenesis of human developmental defects, neuronal disorders, and cancer processes (Fatica and Bozzoni, 2014; Huarte, 2015; Yang et al., 2021).

Human HOTAIR is a 2.2 kb RNA transcribed from the *HOXC* locus, and is expressed in posterior and distal body sites in accordance with its location within the *HOX* locus. HOTAIR binds both Polycomb Repressive Complex 2 (PRC2) and LSD1 complexes and recruits them to multiple genomic sites to promote coordinated H3K27 methylation and H3K4 demethylation, respectively, for gene silencing (Chu et al., 2011; Rinn et al., 2007; Tsai et al., 2010). Although the primary sequences of HOTAIR are poorly conserved between humans and mice, human and murine HOTAIR orthologs demonstrate some functional similarities and conserved RNA secondary structures (Li et al., 2013; Somarowthu et al., 2015). Nonetheless, the sequence divergence creates a challenge to study human HOTAIR *in vivo*.

The overexpression of HOTAIR in several types of human cancers, including breast cancers has been linked to poor survival and increased metastasis (Gupta et al., 2010; Kim et al., 2013; Kogo et al., 2011). Within breast cancer, HOTAIR has been found to be several hundred-fold more highly expressed in metastatic breast tumors and primary breast tumors destined to metastasize (Gupta et al., 2010). HOTAIR overexpression in primary breast cancers is a significant and independent predictor of subsequent metastasis and death across multiple independent cohorts (Gupta et al., 2010; Sorensen et al., 2013). A variety of evidence implicates HOTAIR as a key epigenetic regulator of breast cancer metastasis (Gupta et al., 2010; Liu et al., 2021), but the details of the regulatory mechanism are unclear. Moreover, the mechanism of “epigenetic memory” renders it possible for epigenetic regulators to induce a stable transcriptional program that is retained and propagated by cells without constitutive expression of the epigenetic modulators (Amabile et al., 2016; Bintu et al., 2016; Nunez et al., 2021; Park et al., 2019; Tarjan et al., 2019; Van et al., 2021). For example, the histone methyltransferase Ezh2, a component of the PRC2 complex, can induce long-term silencing by creating a heterochromatin environment at endogenous human gene loci (O’Geen et al., 2019). However, whether HOTAIR-mediated epigenetic and transcriptomic alterations participate in long-term epigenetic memory has not yet been demonstrated. Consequently, it is unclear whether epigenetic changes mediated by HOTAIR can be recovered as a possible avenue of clinical treatment.

In this study, we generate a transgenic murine model with inducible human HOTAIR expression to mechanistically investigate how HOTAIR acts to promote breast cancer progression, and whether depletion of HOTAIR can serve as a therapeutic strategy for treating breast cancer in patients. Using the well-characterized MMTV-PyMT system for endogenous breast cancer development, we confirmed that the induced overexpression of HOTAIR facilitates breast cancer metastasis to the lung and an ongoing dependency on HOTAIR for this phenotype. We also found alterations in both the transcriptome and chromatin accessibility of multiple metastasis associated genes after HOTAIR overexpression and such trend is partially reverted upon HOTAIR silencing, suggesting a continual role of HOTAIR in regulating gene expression by modifying chromatin accessibility.

## Results

### Generation of transgenic mouse with inducible HOTAIR

To better understand the physiological functions of HOTAIR *in vivo*, we generated a transgenic murine model in which human HOTAIR can be conditionally overexpressed upon Doxycycline (Dox) administration. Human HOTAIR cDNA was integrated downstream of a tetracycline response element (TRE) at the *HPRT* locus through inducible cassette exchange recombination in A2Lox.cre mouse embryonic stem cells (mESC) (Iacovino et al., 2011). In this binary control system, the transcription of reverse tetracycline transactivator (rtTA) protein is stimulated by the addition of 1-2 μg/mL of Dox effector, leading to HOTAIR overexpression upon rtTA binding of TRE (Figure 1A). We confirmed that HOTAIR can be effectively induced in mESC with the addition of the drug (Figure S1A). In addition, we validated that upon withdrawing Dox treatment in these ES cells, HOTAIR overexpression returned to baseline levels within 48 hours (Figure 1B). Based on this ES cell line, we then generated a HOTAIR inducible expression mouse model (hereafter named HOTAIR-rtTA mouse) via ES micro-injection and subsequent chimera selection, backcrossing to confirm stable germline transmission of the desired construct. HOTAIR-rtTA mice fed with Dox consistently expressed exponentially higher levels of HOTAIR compared to untreated control mice, as verified through RT-qPCR (Figure 1C).

**Figure 1:**
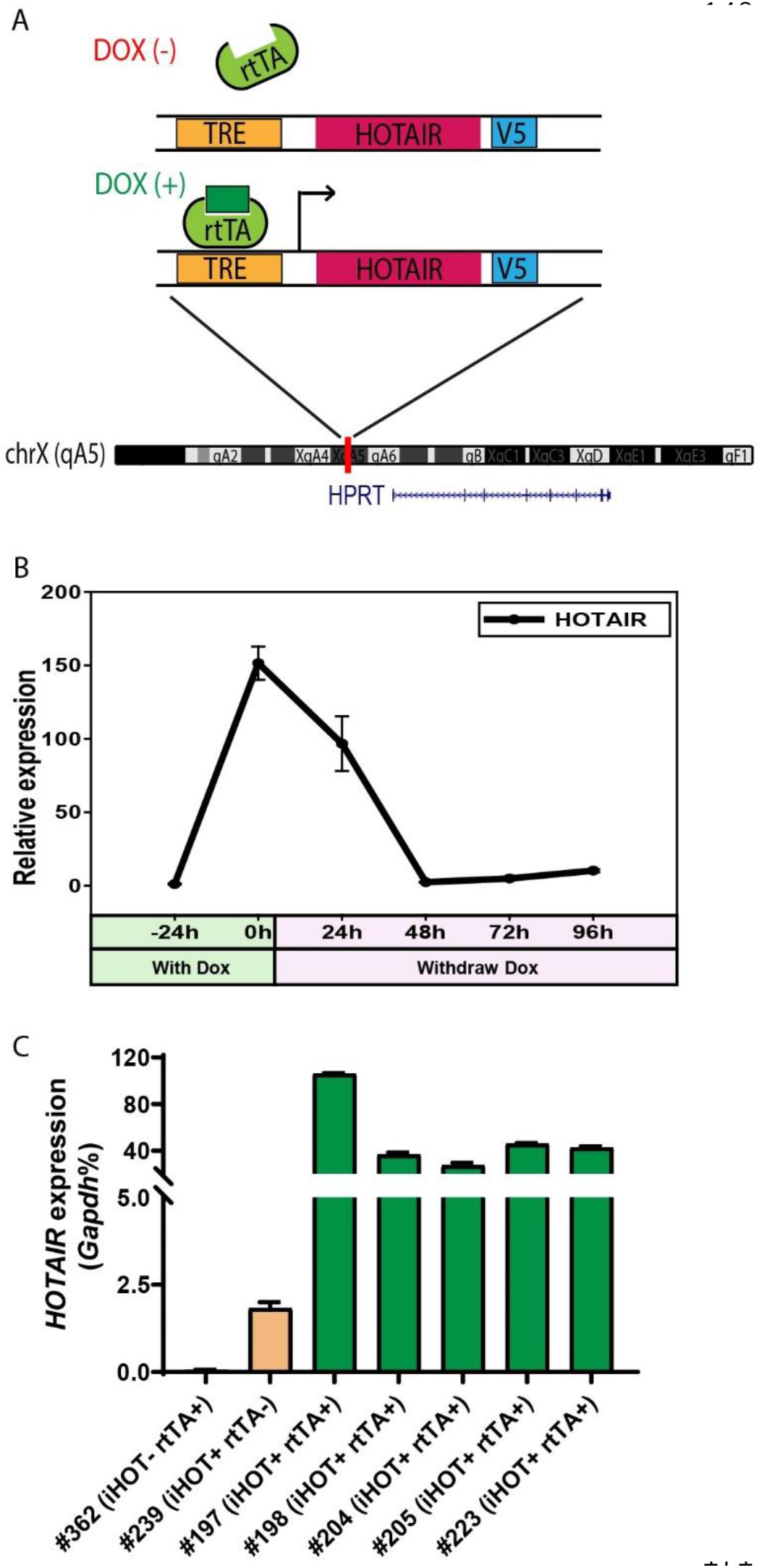
Generation of the inducible HOTAIR murine model. A. Schematic description of the Doxycyclin (Dox) inducible HOTAIR (iHOT) construct and insertion loci. B. *HOTAIR* expression decreased to baseline levels after Dox withdrawal in inducible HOTAIR ES cells. Inducible HOTAIR ES cells were treated with 1 μg/mL Dox for 24 h and then withdrew Dox treatment for 96 h. Human *HOTAIR* expression was detected by qRT-PCR and showed as fold change relative to -24 h. Values are means ± SD. C. High levels of HOTAIR are induced in HOTAIR-rtTA mice (iHOT+ rtTA+) compare to the control group (iHOT ^-^ rtTA ^-^ or iHOT+ rtTA-) after Dox administration. All the mice were treated with Dox for about 10 days. RNA was extracted from mice tails. Human *HOTAIR* levels were quantified by RT-qPCR and calculated as percentages of the mouse *Gapdh* transcript. Values are means ± SD.

In order to investigate whether HOTAIR overexpression is associated with a visible phenotype, mice were fed with Dox water to induce high HOTAIR levels for as long as 1 year. We then evaluated the comprehensive morphology and metabolic features of different tissues. In most cases, we observed little to no change compared to wild type littermates. The subtlety of these phenotypes suggests that HOTAIR overexpression in mice has little effect in a normal genetic background (Figure S1B).

### Elevated HOTAIR promotes breast cancer metastasis to the lung

High levels of HOTAIR are associated with poor survival and more aggressive breast cancers in xenograft models (Gupta et al., 2010). Human breast cancer cells which highly overexpress HOTAIR also display more metastatic and invasive properties compared to control cells (Gupta et al., 2010). To further study the function and mechanism of HOTAIR in promoting breast cancer progression, we engineered the inducible HOTAIR (iHOT) genetic system into a breast cancer model background. We crossed HOTAIR-rtTA mice to MMTV-PyMT (mouse mammary tumor virus-polyoma middle tumor-antigen) mice, a widely utilized breast cancer model in which mice spontaneously develop breast tumors at 8-12 weeks (Figure 2A, hereafter referred to as iHOT-PyMT mouse). This system allowed us to manipulate HOTAIR overexpression using Dox in mice with breast tumors and study HOTAIR function in tumor progression and metastasis in a controlled manner. We confirmed HOTAIR induction in this breast cancer model after Dox treatment using RT-qPCR, and found that HOTAIR transcript expression in PyMT mice containing the iHOT construct (iHOT^+^ Dox^+^ mice) was several hundred-fold higher compared to untreated iHOT mice (iHOT^+^ Dox^-^ mice) or control mice which did not contain the iHOT construct (iHOT^-^ mice) (Figure 2B). We continued to treat the mice with Dox for 3-4 months, exposing mice to continual HOTAIR overexpression, to investigate whether HOTAIR can affect breast tumor progression. After subsequent dissection of lung sections and hematoxylin and eosin staining (H&E staining), we found that primary breast tumor weight and tumor numbers were not affected by continuous exposure to elevated levels of HOTAIR (Figure 2C). In contrast, we observed a greater number of metastatic tumors and larger tumor volumes in the lungs of mice continually exposed to induced HOTAIR (iHOT^+^ mice +Dox treatment, Figure 2D-E, S3A), indicating that HOTAIR overexpression drives metastatic progression. In addition, we also observed that Dox itself has no significant influence in tumor progression of MMTV-PyMT mice as previously reported (Figure S2A) (Nwokafor et al., 2016; Rumney et al., 2017). To reduce the effect of endogenous mouse Hotair, we crossed our iHOT-PyMT mouse model to a murine *Hotair KO* mouse detailed in our previous work (Li et al., 2013). We then selected progeny carrying the appropriate genotypes. We found that expression of endogenous mouse *Hotair* was very low (FigureS2B) and that the knockout of this gene has no significant influence on breast tumor metastasis in MMTV-PyMT mice (Figure S2C). These results demonstrate that high HOTAIR expression promotes breast cancer metastasis in an autochthonous cancer model with intact immune system, consistent with previous clinical observations in patients and studies using xenograft models.

**Figure 2:**
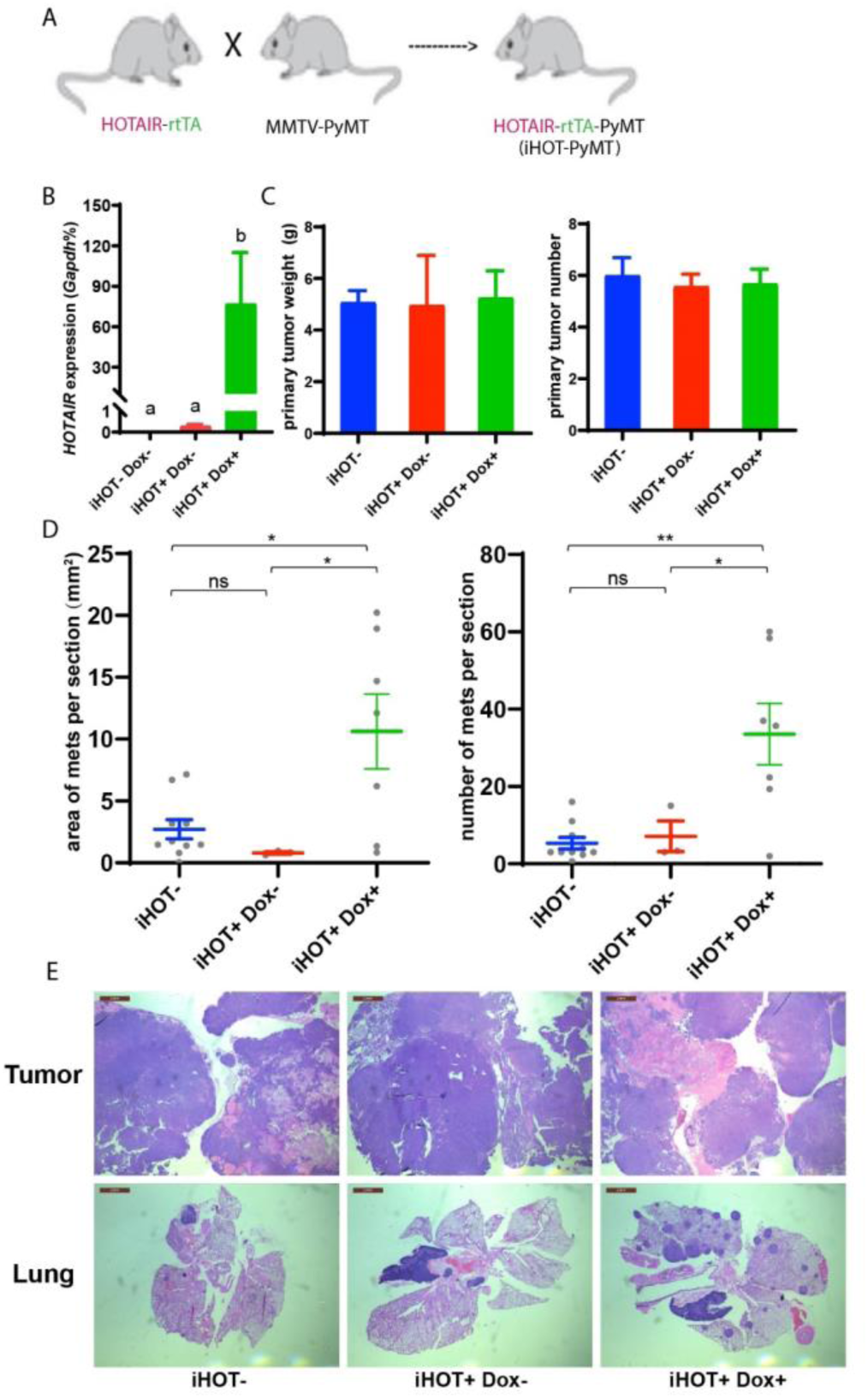
Induced HOTAIR promotes breast cancer metastasis in vivo. A. Cross schema for generating the inducible HOTAIR construct in MMTV-PyMT genetic background. B. qRT-PCR measurements demonstrate that Dox treated iHOT-PyMT mice display significantly higher levels of HOTAIR expression compared to untreated control groups or controls lacking the iHOT construct. Values are means ± SEM, n = 2-3. One-way ANOVA was performed to compare the HOTAIR expression levels between each group (P<0.05). C. There are no statistically significant differences between primary breast tumor mass in grams or number of tumors between Dox+ treated HOTAIR overexpressing mice compared to the untreated control group or controls lacking the inducible HOTAIR construct. Values are means ± SEM, n = 7-9, mice are 3-4 months old. One-way ANOVA was performed between each group and showed no significant differences. D. D-E. iHOT ^+^ mice treated with Dox display an increased number of lung metastases with greater volume. Quantification of lung metastases based on hematoxylin and eosin (H&E) staining of lung and primary tumor sections of iHOT ^+^ Dox treated mice, untreated controls, and controls lacking the iHOT construct. Tumor area in sections were calculated by ImageJ. Dashes represent means ± SEM, n = 3-10. Spots represent every single value. One-way ANOVA was performed (*P<0.05,**P<0.01, ns: not significant). Scale bar = 2 mm.

### Oncogene addiction to HOTAIR in breast cancer cells

As individual mice are variable in tumor progression, we isolated breast cancer cells from the primary tumors of iHOT-PyMT mice (abbreviated as iHOT^+^ cells) in order to gain further mechanistic insight into the functionality of HOTAIR (Figure 3A). In this manner, we can manipulate HOTAIR expression *in vitro* in late stage breast tumor cells in the same genetic background as in our iHOT-PyMT mouse model. We are also able to ascribe the potential impact of HOTAIR as intrinsic to cancer cells. HOTAIR RNA was hundreds-fold higher expressed in iHOT^+^ cells after Dox treatment and levels quickly decreased within 24h after Dox withdrawal, consistent with our observations in the *in vivo* HOTAIR-rtTA mouse (Figure 3B, 1C).

**Figure 3:**
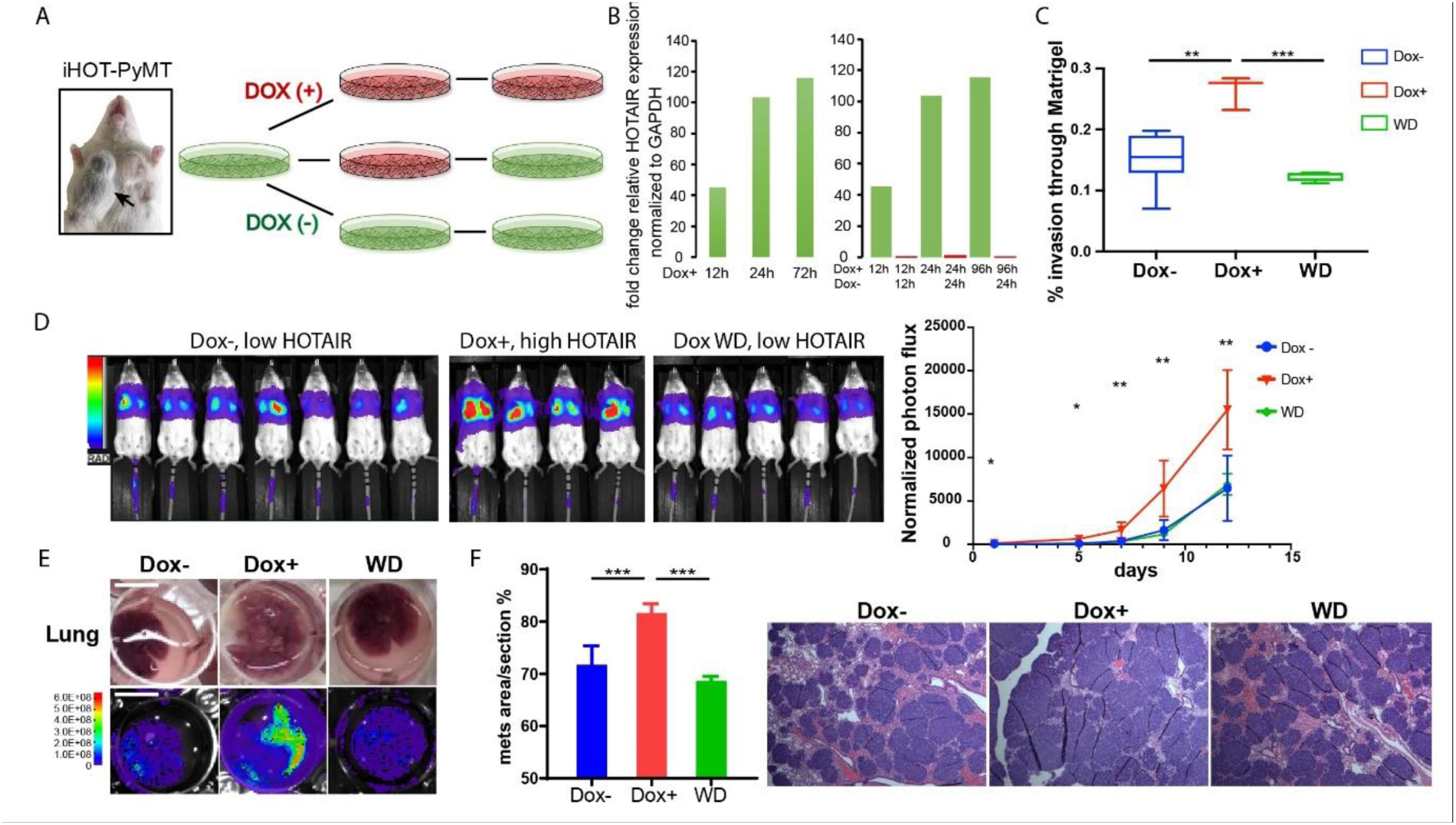
iHOT breast cancer cells display HOTAIR oncogene addiction. A. Treatment strategy of iHOT ^+^ cells to assess the physiological properties of HOTAIR overexpressing cells *in vitro*. iHOT ^+^ cells are isolated from primary mammary tumors of iHOT-PyMT mice. There are three treatment groups: Dox ^-^, Dox ^+^, Dox withdraw after Dox treatment (WD). B. qRT-PCR measurement of HOTAIR expression in iHOT ^+^ cells after Dox treatment for indicated time and withdrawal at 12 or 24 h. HOTAIR is effectively induced after Dox treatment and rapidly decreased after Dox withdrawal in iHOT ^+^ breast cancer cells. C. Dox-induced HOTAIR overexpression promotes increased invasive capacity in iHOT ^+^ cells and the increased invasive capacity of these cells can be rescued when HOTAIR is restored to baseline levels after Dox withdrawal. Matrix invasion assay of iHOT ^+^ breast cancer cells in Dox ^-^, Dox ^+^, Dox withdraw conditions was performed. Invasive capacity was measured by quantifying the displacement of iHOT ^+^ breast cancer cells through Matrigel in media which did or did not contain 2 μg/mL. Values were showed as box plot (n = 3-8). Student t-test was performed between each group, **P<0.01, ***P<0.001. D. Tail vein injection assay performed in SCID mice using iHOT ^+^ breast cancer cells in Dox ^-^, Dox ^+^, Dox withdrawal conditions carrying a luciferase reporter. Dox treated iHOT ^+^ cells overexpressing HOTAIR display higher rates of lung colonization compare to the Dox ^-^ group (n = 4-7, another batch in figure S3D). Student t-test was performed between Dox+ and Dox-group at each time point, *P<0.05, **P<0.05. We show that the potential for lung colonization by iHOT ^+^ cells decreases after Dox withdrawal. E. Representative photos and bioluminescent imaging of lungs dissected from SCID mice at the 2 week timepoint after tail vein injection of iHOT ^+^ cells. Scale bar = 1 cm. F. Quantified percentage of metastatic lung tumor area in Dox ^-^, Dox ^+^, Dox withdrawal cells after tail vein injection in lung sections with HE staining. Values are means ± SD, n=4-7. One-way ANOVA was performed between each group, ***P<0.001.

We next examined the effects of manipulating HOTAIR level in iHOT^+^ murine breast cancer cell lines *in vitro*. To assess the metastatic capacity of HOTAIR overexpressing cells as compared to control cells, we induced HOTAIR overexpression in iHOT^+^ breast cancer cells by Dox treatment for 7-18 days and subsequently performed a Matrigel invasion assay, which measures the ability of cells to migrate through a basement-membrane like extracellular matrix (Figure 3C). We found that HOTAIR overexpression promotes the invasive capacity of breast cancer cells. In order to test whether HOTAIR overexpression evokes epigenetic “memory” resulting in a phenotype of increased invasiveness, we performed a complete Dox withdrawal for 7-8 weeks. We observed that cellular invasion decreased after HOTAIR overexpressing cells underwent complete Dox withdrawal, suggesting that ongoing HOTAIR is required to promote increased cancer cell metastasis (Figure 3C). We also measured the growth curve of iHOT^+^ cells under these same three conditions and found that HOTAIR expression levels do not affect proliferation of iHOT^+^ breast cancer cells (Figure S3C).

To study the impact of HOTAIR overexpression on the metastatic potential of iHOT^+^ cells *in vivo*, we labelled iHOT^+^ cells with the firefly luciferase reporter gene *luc2*, enabling sensitive bioluminescence imaging of HOTAIR expression in live animals. We injected *luc2-*labeled iHOT^+^ cells exposed to Dox for 18 days, iHOT^+^ cells treated with Dox for 10 days and subsequently withdrawn from Dox, and untreated iHOT^+^ control cells through the tail vein of female SCID Beige mice. iHOT^+^ overexpression was sustained in live animals by feeding the experimental group Dox water over time. We then imaged live animals for luciferase bioluminescence every 2-3 days for around 2 weeks and measured the rates of lung colonization by injected iHOT^+^ tumor cells. As expected, mice injected with Dox treated iHOT^+^ cells displayed significantly higher lung colonization after tail vein injection compared untreated control groups. Mice injected with cells that had been treated with Dox and subsequently withdrawn displayed low lung colonization of fluorescent cells, similar to the bioluminescence profile of mice injected untreated iHOT^+^ control cells (Figure 3D&E, Figure S3D). Altogether, our results suggest that the increased metastatic potential of HOTAIR overexpressing breast cancer cells requires ongoing HOTAIR activity and is abrogated by silencing HOTAIR.

We sacrificed injected mice at the two-week time point and dissected lung tissue from the mice. We performed histologic analysis of lung sections using hematoxylin and eosin staining to visualize tumor metastasis. The lung sections of mice injected with HOTAIR overexpressing cells displayed a higher metastatic tumor burden compared to mice injected with HOTAIR withdrawn cells or untreated control cells (Figure 3E&F). In contrast, mice injected with HOTAIR withdrawn cells where HOTAIR expression level is returned to baseline display similar levels of lung colonization compared to injection with untreated control cells. In brief, we conclude that breast cancer cells with elevated HOTAIR require persistent and sustained HOTAIR overexpression to retain a highly invasive and metastatic phenotype.

### HOTAIR is required for metastasis-associated chromatin reprogramming and gene expression

We hypothesize that HOTAIR dysregulation during breast cancer metastasis leads to an altered chromatin landscape and subsequent changes in gene expression in affected cells by recruiting chromatin remodeling machinery such as the Polycomb complex (PRC2). PRC2 promotes chromatin compaction through catalyzing the trimethylation of histone H3 at lysine 27 (H3K27me3), a repressive histone mark. To this end, we analyzed gene expression patterns in iHOT^+^ cells treated with 3 conditions: Dox^+^ HOTAIR overexpressing cells which were consistently treated with Dox for several days; Dox^WD^ cells which had been temporarily treated with Dox and later withdrawn from HOTAIR overexpression; and untreated control cells in which HOTAIR remained at baseline levels. We performed RNA-seq to gain insight into the molecular basis of the observed phenotypes within these experimental conditions. First, we checked HOTAIR expression levels in the RNA-seq data, which revealed that HOTAIR levels were elevated in the Dox^+^ group and reverted to basal level in the Dox^WD^ group (Figure S3B). RNA-seq analysis revealed that there were 219 up-regulated genes and 280 down-regulated genes in the Dox treated HOTAIR overexpressing group compared to the untreated control group (FDR < 0.05, fold change > 2, Figure 4A).

**Figure 4:**
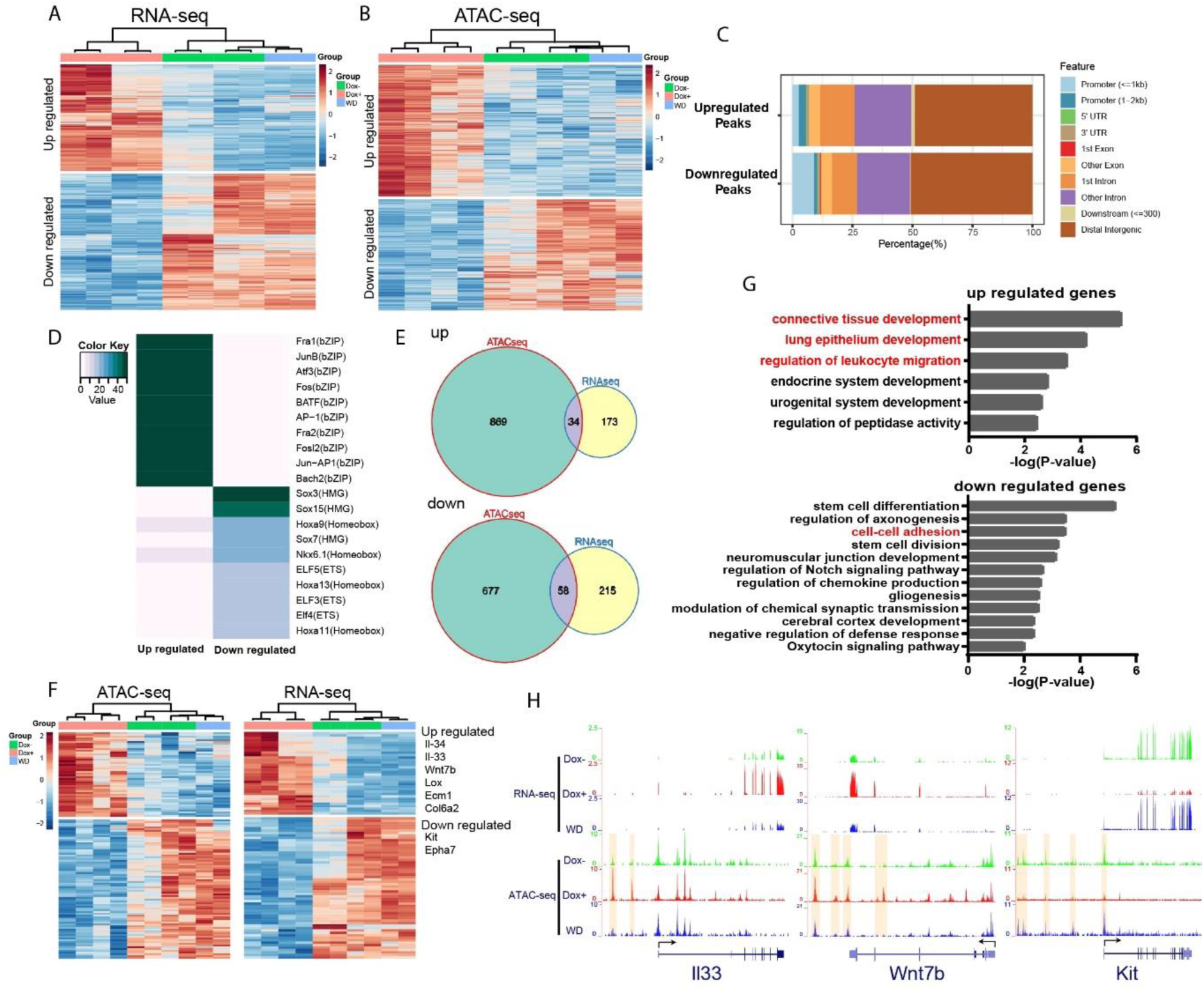
HOTAIR is required for metastasis-associated chromatin accessibility and genes expression. A: Hierarchical clustering heatmap of RNA-seq data of iHOT ^+^ cells subject to either Dox ^-^, Dox ^+^, or Dox withdrawal treatment. Dox induced HOTAIR overexpression results in altered gene expression profiles. There are 499 genes differentially expressed between Dox ^+^ and Dox ^-^ cells (FDR < 0.05, Fold change >2). Transcriptome patterns of Dox withdrawal cells are similar to Dox ^-^ cells. B: Hierarchical clustering heatmap of differential ATAC-seq peaks of iHOT ^+^ cells with Dox ^-^, Dox ^+^, Dox withdraw treatments. Dox induced HOTAIR overexpression alters chromatin accessibility. There are 1933 differential peaks between Dox+ and Dox-cells. Chromatin accessibility landscape of differential peaks returns to Dox ^-^ status after Dox withdrawal. C: Feature distribution of differentially regulated peaks. We observe significantly more downregulated peaks located in promoter regions (less than 1 kb to transcription start sites) compared to upregulated peaks (p<0.01). D: Enriched motifs of differential ATAC-Seq peaks by HOMER motif analysis. Color coding indicates -log10 p values. E: Venn diagram representing overlap between ATAC-Seq differential peak-associated genes and differentially expressed genes by RNA-Seq (up: up-regulated genes; down: down-regulated genes). F: Heatmap of transcriptome and corresponding chromatin status of overlapping genes in 4E in Dox ^-^, Dox ^+^, and Dox withdrawal conditions. Listed on the right are several representative genes related to cancer cell metastasis. G: Gene ontology of overlapping genes in Fig. 4E. Terms highlighted in red are examples of terms related to cancer cell metastasis. H: HOTAIR regulates chromatin accessibility and gene expression. Normalized ATAC-Seq and RNA-Seq sequencing tracks of metastasis-related genes *Il33, Wnt7b* and *Kit*. Alteration of chromatin accessibility correlates well with transcriptional changes.

We then applied a GO term analysis to these groups of differentially expressed genes and found terms consistent with invasion and metastasis phenotypes (Figure S4A). We discovered that GO terms such as “epithelial cell proliferation” and “Wnt Signaling Pathway and Pluripotency” were enriched in the up-regulated gene group, and terms such as “regulation of cell adhesion” were enriched in the down-regulated gene group. Importantly, in the Dox withdrawal condition, we observed that most of these differentially expressed genes to revert to expression levels similar to that of the untreated control group. This result suggests that genes that are differentially expressed due to high levels of HOTAIR transcript in the system can be subsequently reversed to their original expression level upon HOTAIR withdrawal (Figure 4A). We visualized gene expression changes between experimental groups using a hierarchical clustering heatmap, revealing that the transcriptome patterns of Dox withdrawal cells are similar to Dox^-^ cells (Figure 4A). This supports our previous *in vivo* observations, which imply that the increased invasiveness and metastatic properties of breast cancer cells can be reversed with HOTAIR withdrawal.

Since HOTAIR is known to reprogram chromatin status and alter chromatin modifications to promote cancer metastasis (Gupta et al., 2010), we performed an Assay for Transposase-Accessible Chromatin using sequencing (ATAC-Seq) to assess genome-wide changes in the chromatin landscape induced by HOTAIR overexpression. High levels of HOTAIR induced by Dox treatment led to altered chromatin accessibility (Figure 4B). There are in total 1933 peaks with differential accessibility between all HOTAIR high Dox+ and all HOTAIR low Dox-cells, of which 1048 peaks are up-regulated and 886 peaks are down-regulated. We observed both up-regulated and down-regulated peaks to be mainly located in distal intergenic regions (about 50%). We found that a higher percentage of down-regulated peaks occur in promoter regions less than 1 kb, indicating that HOTAIR most likely functions suppressively near promoter regions to modulate the expression of down-regulated genes (Figure 4C).

To reveal possible functional pathways which HOTAIR participates in within the cell, we performed motif enrichment analysis by HOMER on the up-regulated and down-regulated peaks which we identified through ATAC-seq (Figure 4D). The top motif enriched in up-regulated peaks involved many transcription factors associated with tumor metastasis, such as AP-1 family members Fra1, JunB, Atf3, and Fos (Belguise et al., 2005; Hyakusoku et al., 2016; Milde-Langosch et al., 2004; Yuan et al., 2013). In the top motif enriched in down regulated peaks, Sox7 was reported to play inhibitory roles related to cellular proliferation, migration and invasion in breast cancer (Stovall et al., 2013). Previous work in pancreatic ductal adenocarcinoma has demonstrated that Sox15 is a potential tumor suppressor of the Wnt/β-catenin signaling pathway (Thu et al., 2014). These observations support our primary hypothesis that high levels of HOTAIR transcript promote breast cancer metastasis and suggest that HOTAIR might have functional relevance to those identified factors. We then analyzed peaks associated with known genes and found that cell samples subjected to Dox withdrawal also clustered together with Dox-untreated control samples in a hierarchical clustering heatmap (Figure 4B), consistent with transcriptomic patterns which we observed in previous experiments. GO analysis of these peaks associated with known genes showed enrichment of many terms related to cell adhesion (Figure S4B). For example, the terms “regulation of cell adhesion” and “leukocyte migration” were enriched in up-regulated peaks associated with genes, whereas terms such as “cell-cell adhesion”, “cell-substrate adhesion” and “negative regulation of locomotion” were enriched in down-regulated peaks associated with genes (Figure S4B). Consistent with our previous results in our RNA-seq experiments, the chromatin accessibility level of most differential peaks identified with ATAC-seq can be subsequently rescued by HOTAIR withdrawal in the Dox^WD^ HOTAIR low group (Figure 4B).

To further investigate downstream pathway components regulated by HOTAIR, we compared differentially accessible peaks in the immediate vicinity of differentially expressed genes, and from the overlapping set identified 34 up-regulated genes and 56 down-regulated genes associated with peaks. Many of the selected genes were linked to metastasis, such as *Il-33, Il-34, Wnt7b, Lox, Ecm1, Col6a2* (up regulated), *Kit*, and *Epha7* (down regulated) (Figure 4E&F). GO term analysis revealed these up-regulated genes are associated with pathways such as “connective tissue development”, “lung epithelium development” and “regulation of leukocyte migration”, while down-regulated genes are associated with “cell-cell adhesion”, all of which are metastasis-related cellular processes (Figure 4G). As shown in examples from Figure 4H, both *II33* and *Wnt7b* were up-regulated with HOTAIR induction (Dox^+^) and reverted to low expression levels after HOTAIR withdrawal (WD). Conversely, the transcription factor Kit was down-regulated after HOTAIR induction and reverted to high expression levels after HOTAIR withdrawal.

Several lines of evidence in the literature support the interpretation that our model accurately captures molecular changes corresponding to cancer progression and the development of metastatic cellular phenotypes. For example, ATAC-seq signals in the regulation region of the differentially expressed genes we identified display a consistent change in the ligand (Figure 4H) IL-33 and its receptor ST2, which form the IL-33 /ST2 signaling pathway. This pathway has previously been linked with tumor progression, cellular invasion, and metastasis (Kim et al., 2015; Larsen et al., 2018; Zhang et al., 2019). The WNT/β-catenin pathway plays a critical role in tumorigenesis, and Wnt7b was reported to function in mediating metastasis in breast cancer (Xu et al., 2020; Yeo et al., 2014). Kit, also known as c-Kit, CD117 (cluster of differentiation 117) or SCFR (mast/stem cell growth factor receptor), is one of the receptor tyrosine kinases (RTK), and has been previously reported to be involved in tumour migration or invasion (Golubovskaya and Wu, 2016). Altogether, these results suggest that HOTAIR transcript regulates the expression of multiple metastasis-associated genes by modifying their chromatin accessibility to promote tumour metastasis.

We observed that some adhesion-associated GO terms were enriched for HOTAIR-regulated genes and that the expression level of several EMT hallmark genes such as *Lox, Ecm1, Col6a2* were significantly altered with HOTAIR overexpression (Figure 4G&F). To investigate the metastatic phenotype of iHOT^+^ cells, we performed immunofluorescence staining of prototypical EMT markers. After Dox treatment, the mesenchymal marker Vimentin was induced and the epithelial marker E-cadherin was reduced. In addition, the iHOT^+^ cells also displayed more spindle-like morphology and cell detachment from epithelioid colonies (Figure 5A&B). When Dox was withdrawn for a brief period over 3 cellular passages, Vimentin was still expressed albeit at a reduced level. However, when Dox was withdrawn for an extended period of time over 10 cellular passages, Vimentin was largely abrogated and E-cadherin re-expressed to similar levels as in untreated cells, and Dox withdrawn cells returned to epithelial morphology (Figure 5A&B). These results demonstrate that iHOT^+^ cells undergo EMT when exposed to HOTAIR overexpression as a result of treatment with dox, and recover their original epithelial states after longer periods of withdrawal from Dox, which is consistent with the invasion and metastasis phenotypes we previously observed. However, it appears that effects modulated by HOTAIR might persist for some time. For example, as HOTAIR expression is quickly restored to basal levels within 24 h after Dox withdrawal (Figure 3B), elevated Vimentin expression endures over several days lasting as long as 3 passages (Figure 5B).

**Figure 5:**
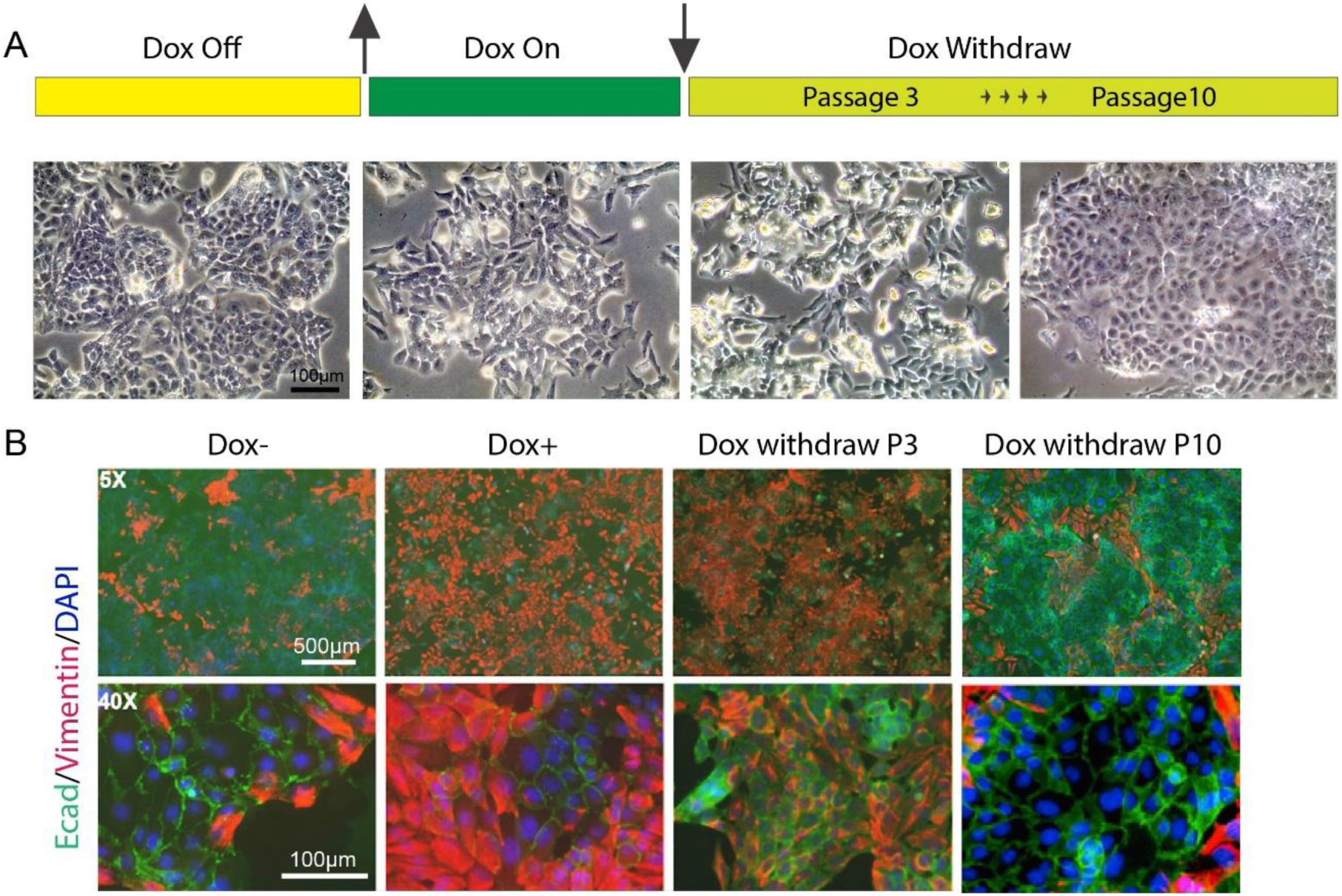
HOTAIR continually enforces epithelial-mesenchymal transition of breast cancer cells. A. Scheme demonstrating how iHOT ^+^ cells were treated with Dox and microphotographs of cell morphology in each condition. B: Immunofluorescence staining of E-cadherin (green), Vimentin (red) and DAPI (blue) of iHOT ^+^ cells in each condition.

Collectively, our results demonstrate that the cellular function of HOTAIR as a regulator of gene expression is conserved between mammalian lineages and confirm that exposure to sustained high levels of HOTAIR over time facilitates increased EMT, invasiveness and metastatic capacity in breast cancer cells. We also observe alterations in both the cellular transcriptome as well as the chromatin accessibility landscape of multiple metastasis-associated genes after HOTAIR overexpression. Subsequently silencing HOTAIR expression for enough time abrogates this metastatic phenotype, as well as any resulting alterations in transcriptome and chromatin accessibility, indicating that these HOTAIR-modulated alterations have not yet formed persistent epigenetic memory. Thus, HOTAIR is required in an ongoing manner to reprogram oncogene expression. Our results also suggest that targeting overexpressed HOTAIR transcript can potentially serve as a therapeutic strategy in breast cancer patients to limit metastatic progression.

## Discussion

Overexpression of the lncRNA HOTAIR is significantly correlated with tumor progression and poor prognosis in many kinds of tumors and is a powerful predictor of tumor metastasis. As such, targeting HOTAIR overexpression is of clinical interest as a potential therapy for cancer patients (Gupta et al., 2010; Mozdarani et al., 2020). Hence, there exists a critical need to systematically interrogate and understand the physiological functions and mechanisms of action of HOTAIR in model organisms. The HOTAIR-rtTA-PYMT mouse generated in this paper provides a model to analyze the functional role of human HOTAIR *in vivo* within the context of breast cancer metastasis.

Murine HOTAIR and human HOTAIR transcripts are conserved in structure rather than primary sequence (He et al., 2011; Yu et al., 2012). It was previously reported that human HOTAIR binds to both the PRC2 and LSD1 chromatin remodeling complexes, acting as a scaffold to target these complexes to the HOX loci and enforcing a silenced chromatin state at this position (Tsai et al., 2010). Human HOTAIR may have additional gene silencing mechanisms (Meredith et al., 2016). Murine HOTAIR has been observed to function as a trans-acting regulator similar to human HOTAIR, capable of acting *in trans* to influence the activities of distal genes to its site of origin (Li et al., 2013). However, there exists no direct evidence of this functional conservation. In our inducible human HOTAIR murine model, human HOTAIR transcript overexpression was found to promote breast cancer metastasis *in vivo*, consistent with previous observations in patients and human breast cancer cell lines. Based on our ATAC-seq data, we find significant changes in the chromatin landscape when human HOTAIR is upregulated in mouse tissue, indicating that human HOTAIR functions through a similar mechanism as murine HOTAIR by binding chromatin remodeling machinery and targeting the biochemical activities of these complexes to specific genomic loci. We postulate that this reprogramming of the underlying chromatin by HOTAIR leads to altered cellular function, which allows the cell to metastasize and thrive at ectopic sites.

Many lncRNAs function in regulating epigenetic modifications, some of which are also involved in the establishment of persistent epigenetic memory (Arunkumar et al., 2022; Begolli et al., 2019; Fok et al., 2019; Wei et al., 2017). For example, recent work showed that transcription of locus 8q24-derived oncogenic lncRNAs such as *PCAT2* could recruit centromeric protein-A (CENP-A), a variant of Histone H3, promote the ectopic localization of CENP-A; and alter epigenetic memory at a fragile chromosomal site in human cancer cells (Arunkumar et al., 2022). In the current study, we investigated whether HOTAIR-mediated epigenetic alterations participate in enacting long-term epigenetic memory using transgenic iHOT^+^ tumor cells. Our results demonstrate that a complete Dox withdrawal could restore all alterations in either transcriptomes, chromatin states or metastatic phenotypes. Our data implies that the presence of HOTAIR transcript is required to promote cancer metastasis and propagate changes in the chromatin landscape. This indicates that HOTAIR is needed to enforce cellular transcriptional programs—a “lncRNA addiction”. In contrast, Xist lncRNA is involved in establishment of X chromosome inactivation in female cells but is largely dispensable for the maintenance of the inactive state over subsequent cell divisions and embryonic development (Yu et al., 2021). In this regard, the role of HOTAIR is more akin to a transcription factor that mediates a large-scale but reversible program of gene expression. We demonstrated that restoring HOTAIR expression level to baseline from a highly overexpressed state in iHOT^+^ breast cancer cells isolated from the primary tumors of iHOT-PyMT mice could rescue expression of most altered genes and chromatin accessibility (Figure 4A&B), as well as restoring the overall physical phenotype of the organism (Figure 3C-F&Figure 5). Our results suggest that persistent and sustained HOTAIR overexpression engender high metastatic potential within cancerous cells, further implying HOTAIR’s utility as a therapeutic target.

Our mouse model of human HOTAIR function provides an in vivo system to test small molecules or antisense oligonucleotides as potential therapeutics in the future. LncRNA can be targeted by multiple approaches: 1) siRNAs or antisense oligonucleotides (ASOs) can be used to silence genes in a sequence-specific manner to decrease lncRNA levels post-transcriptionally; 2) Modulation of lncRNA expression by the CRISPR/Cas9 System; 3) Inhibition of RNA-protein interactions or prevention secondary structure formation (Arun et al., 2018). Several studies demonstrated that knockdown of HOTAIR can be achieved using siRNA or ASO strategies *in vitro* (Lennox and Behlke, 2016; Rinn et al., 2007; Tsai et al., 2010; Yang et al., 2011), but targeting lncRNAs *in vivo* continues to present significant challenges. A peptide nucleic acid (PNA)-based approach was reported to block the EZH2 binding domain of HOTAIR, inhibiting HOTAIR-EZH2 activity and subsequently decreasing invasion of ovarian and breast cancer cells and resensitize resistant ovarian tumors to platinum (Ozes et al., 2017).

In addition, several drugs have been recognized to indirectly down-regulate HOTAIR expression (Mozdarani et al., 2020). For example, the isoflavone-based drugs Calycosin and Genistein inhibit HOTAIR expression by repressing Akt pathway upstream of HOTAIR transcription (Chen et al., 2015; Ozes et al., 2017). Direct or indirect suppression of the WNT pathway was also reported to down regulate HOTAIR expression in cancer cells (Carrion et al., 2014; Wang et al., 2015). Using a combination of RNA-seq and ATAC-seq assays, we identified a candidate gene set with altered expression or chromatin status in the context of HOTAIR overexpression, which were rescued when HOTAIR expression level was restored. This gene set encompasses the direct or indirect molecular targets of HOTAIR. Future work to delineate the direct targets and cofactors of HOTAIR within these gene set will provide further insight into the molecular mechanism and biological function of the HOTAIR transcript.

## Methods

### Animals

#### Generation of the inducible HOTAIR mESC line

To generate the inducible HOTAIR mESC line, we first cloned human HOTAIR cDNA into p2Lox plasmid (p2Lox-hHOTAIR), then transfected the plasmid into A2Lox.cre mESC lines. The stably integrated cell line was selected through the inducible cassette exchange method, described with details in Iacovino, et al, 2011, Stem Cells (Iacovino et al., 2011).

#### Generation of HOTAIR-rtTA mouse

To generate the inducible HOTAIR transgenic mouse model, we performed micro-injection of the ES cells into the mouse blastocyst, which was allowed to develop into the chimeric animal. After backcrossing to pure C57/BL6 mice and performing genotyping selection, we confirmed that the transgene was stably integrated and germline transmittable animals were picked-up and maintained for the subsequent study.

#### Generation of iHOT-PyMT mouse

To generate a breast cancer animal model with conditional expression of human HOTAIR, we crossed the HOTAIR-rtTA line with a murine breast cancer mouse line MMTV-pyMT (FVB/N-Tg (MMTV-PyVT) 634Mul/J #2374, Jackson Laboratory). Through several rounds of crossing and genotyping, the triple-positive mice (i.e. HOTAIR+, rtTA+, MMTV+) were selected for further study.

All mice were bred in the Stanford University Research Animal Facility in accordance with the guidelines.

Dox treated mice were fed with doxycycline at the concentrations of 5 g/L supplemented with 50 g/L sucrose via drinking water.

### Tumor cell isolation and DOX treatments

iHOT^+^ cells were derived from the primary tumors of iHOT-PyMT mice as described in the following steps. The primary tumors of iHOT-PyMT mice were dissected and chopped into small pieces. The tissues were then digested using 5 Ml collagenase buffer (0.5% collagenase I in HBSS) per gram for 1-1.5 h at 37°C with moderate shaking (∼200 rpm). The suspended cell mixture was then spun at 600 rcf for 2-10 minutes. The pellet of small epithelial sheet was resuspended in 0.25% Trypsin for 10 min and neutralized with 10% FBS/DMEM and 1X DNase 10 min at 37°C. The dissociated cells were passed through a 0.45 μm cell strainer, and the cells were pelleted at 450 rcf for 5min. The cells were then washed with PBS and pelleted again. The cells were then plated and cultured in DMEM/F-12 media with 5% Tet System Approved FBS, 1% P/S Penicillin-Streptomycin, 5 μg/mL insulin from bovine and 1 μg/mL hydrocortisol.

Three conditions of iHOT^+^ cells were used in this study: Dox^+^, Dox^-^ and Dox withdrawal (Dox^WD^). In the Dox^+^ group, iHOT^+^ cells were treated with 2 μg/mL DOX for 7 days or 18 days; in the Dox^-^ group, iHOT^+^ cells were treated with solvent control for 18 days or 59 days; and in the Dox^WD^ group, iHOT^+^ cells were first treated with DOX for 10 days then removed for another 49 days.

### qRT-PCR

Total RNA from mouse tails or iHOT^+^ cells was extracted using TRIzol and the RNeasy mini kit (Qiagen). RNA levels (starting with 50–100 ng per reaction) for a specific gene were measured using the Brilliant SYBR Green II qRT–PCR kit (Strategene) according to manufacturer instructions. All samples were normalized to mouse GAPDH.

Primers used:

hHOTAIR-F: GGTAGAAAAAGCAACCACGAAGC

hHOTAIR-R: ACATAAACCTCTGTCTGTGAGTGCC

mHotair-F: CCTTATAAGCTCATCGGAGCA

mHotair-R: CATTTCTGGGTGGTTCCTTT

mGapdh-F: CTGGAGAAACCTGCCAAGTA

mGapdh-R: TGTTGCTGTAGCCGTATTCA

### Invasion assay

The matrigel invasion assay was done using the Biocoat Matrigel Invasion Chamber as previously described (Gupta et al., 2010). In brief, 5 × 10^4^ cells were plated in the upper chamber in serum-free media. The bottom chamber contained DMEM media with 10% FBS. After 24–48 h, the bottom of the chamber insert was fixed and stained with Diff-Quick stain. Cells on the stained membrane were counted under a dissecting microscope. Each membrane was divided into four quadrants and an average from all four quadrants was calculated.

### Mice tail vein injection

Mouse tail vein xenografts assay was performed as previously described (Gupta et al., 2010). Female athymic nude mice were used. 2.5-3 million iHOT^+^ cells with different Dox treatments in 0.2 ml PBS were injected by the tail vein into individual mice (5-7 for each treatment). Mice were observed generally for signs of illness weekly for the length of the experiment. The lungs were excised and imaged, then fixed in formalin overnight and embedded in paraffin, from which sections were made and stained with haematoxylin and eosin by our pathology consultation service. These slides were examined for the micro-metastases, which were counted in three fields per specimen.

### Bioluminescence imaging

Mice received luciferin (300 mg/kg, 10 min before imaging) and were anaesthetized (3% isoflurane) and imaged in an IVIS spectrum imaging system (Xenogen, part of Caliper Life Sciences). Images were analyzed with Living Image software (Xenogen, part of Caliper Life Sciences). Bioluminescent flux (photons s^-1^ sr^-1^ cm^-2^) was determined for the lungs.

### Immunofluorescence

Cells were fixed using 4% paraformaldehyde solution for 20 min at room temperature. The cells were then permeabilized and blocked in blocking solution (0.1% Triton X-100 and 0.1% BSA in PBS) for 45 min. Afterward, the cells were incubated with anti-Vimentin (ab92547, abcam) and anti-E-cadherin (14-3249, ebioscience) antibodies at 4 °C for 1 h, followed by Alexa Fluor 546-labeled and 488-labeled secondary antibody for 1 h, and counterstained with DAPI.

### RNA-Seq

Total RNA of iHOT^+^ cells with different treatments were extracted by Trizol following RNA clean-5 kit. Poly-A-selected RNA was isolated and the libraties were prepared with the dUTP protocol and sequenced using the Illumina Genome Analyzer IIX platform with 36 bp reads. Raw reads were aligned to the mouse reference sequences NCBI Build 37/mm9 with STAR aligner. Expression levels of RefSeq annotated genes were calculated using RSEM. The differential analysis was conducted using R package DESeq2. The Benjamini hochberg procedure was used to adjust for multiple hypothesis testing. Genes with FDR < 0.05 and fold change of 2 were considered as significant.

### ATAC-Seq

ATAC–seq of iHOT^+^ cells with different treatments were performed as previously described (Corces et al., 2017). Briefly, 50000 cells were pelleted, resuspended in 50 μl of lysis buffer (10 mM Tris-HCl, pH 7.4, 3 mM MgCl_2_, 10 mM NaCl, 0.1% NP-40 (Igepal CA-630)), and immediately centrifuged at 500g for 10 min at 4 °C. The nuclei pellets were resuspended in 50 μl of transposition buffer (25 μl of 2× TD buffer, 22.5 μl of distilled water, 2.5 μl of Illumina Tn5 transposase) and incubated at 37 °C for 30 min. Transposed DNA was purified with the MinElute PCR Purification kit (Qiagen), and eluted in 10 μl of EB buffer.

Adaptor sequence trimming using in-house software and mapping to hg19 using Bowtie2 were performed. The reads were then filtered for mitochondrial reads, low-quality reads, PCR duplicates, and black list regions. The filtered reads for each group were merged, and peak calling was performed by MACS2. The reads in peaks for each individual sample were quantified using BEDTools multicov with the merged MACS2 narrow peaks. Peak counts were then combined into an N × M matrix where N represents called peaks, M represents the samples, and each value Di,j represents the peak intensity for respective peak i in sample j. Each peak was annotated by the nearest gene. This matrix was then normalized using R package DESeq2, and differential analysis was conducted using negative binomial models. The Benjamini hochberg procedure was used to adjust for multiple hypothesis testing. Peaks with FDR < 0.05 and fold change of 2 were considered as significant.

## Authors’ contributions

QM, LL and HYC conceived the project. QM, LL, KT, YZ, UML, QS, M-CT, J-AC, IL and HZ performed the experiments. QM, LY, YZ, LL and KT performed data analysis. QM, LY, KT and HYC wrote the manuscript with inputs from all authors and LZ assisted in manuscript collation. QM, LL and HYC obtained funds.

---

**Supp 1:**
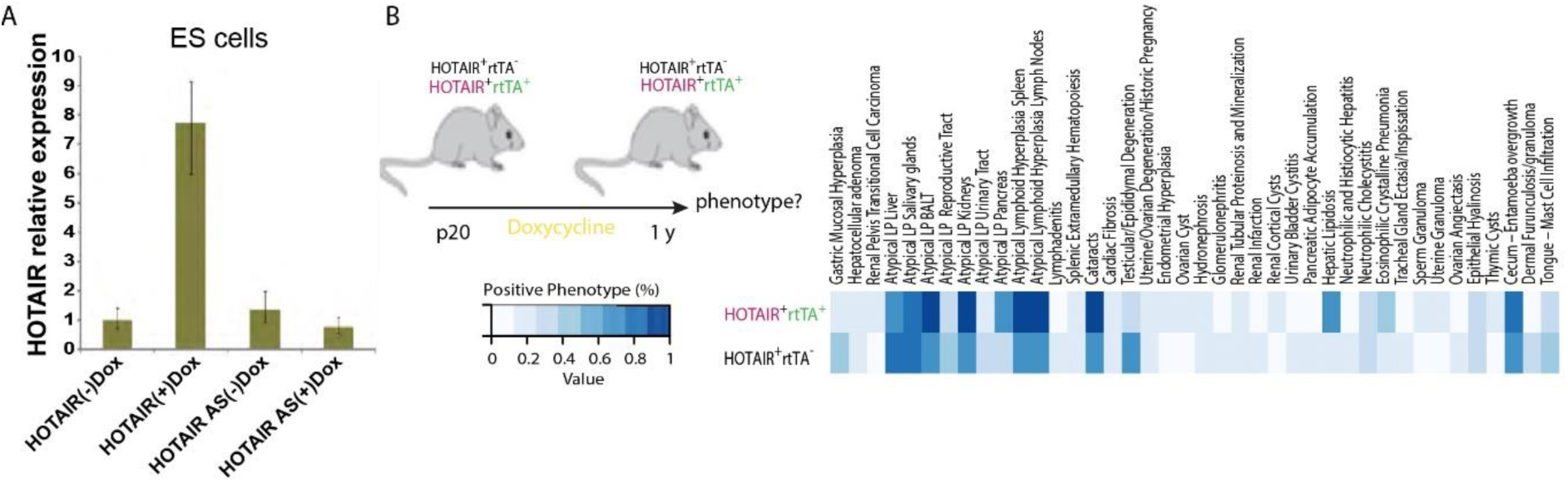
HOTAIR expression can be induced and inducible HOTAIR mice have no obvious phenotypes in different tissues. A: HOTAIR expression in inducible HOTAIR embryonic stem (ES) cells. (AS: HOTAIR antisense controls) B: Inducible HOTAIR mice have no obvious phenotypes in different tissues. 8 weeks old mice were fed with Dox water for 1 year. Phenotypes about hyperplasia and neoplasia, lymphoid system, age-related lesions, urinary system, liver and pancreas, respiratory, reproductive and so on are tested and compared between HOTAIR^+^ rtTA^+^ and HOTAIR^+^ rtTA^+^ mouse. Each genotype has 6 mice.

**Supp 2:**
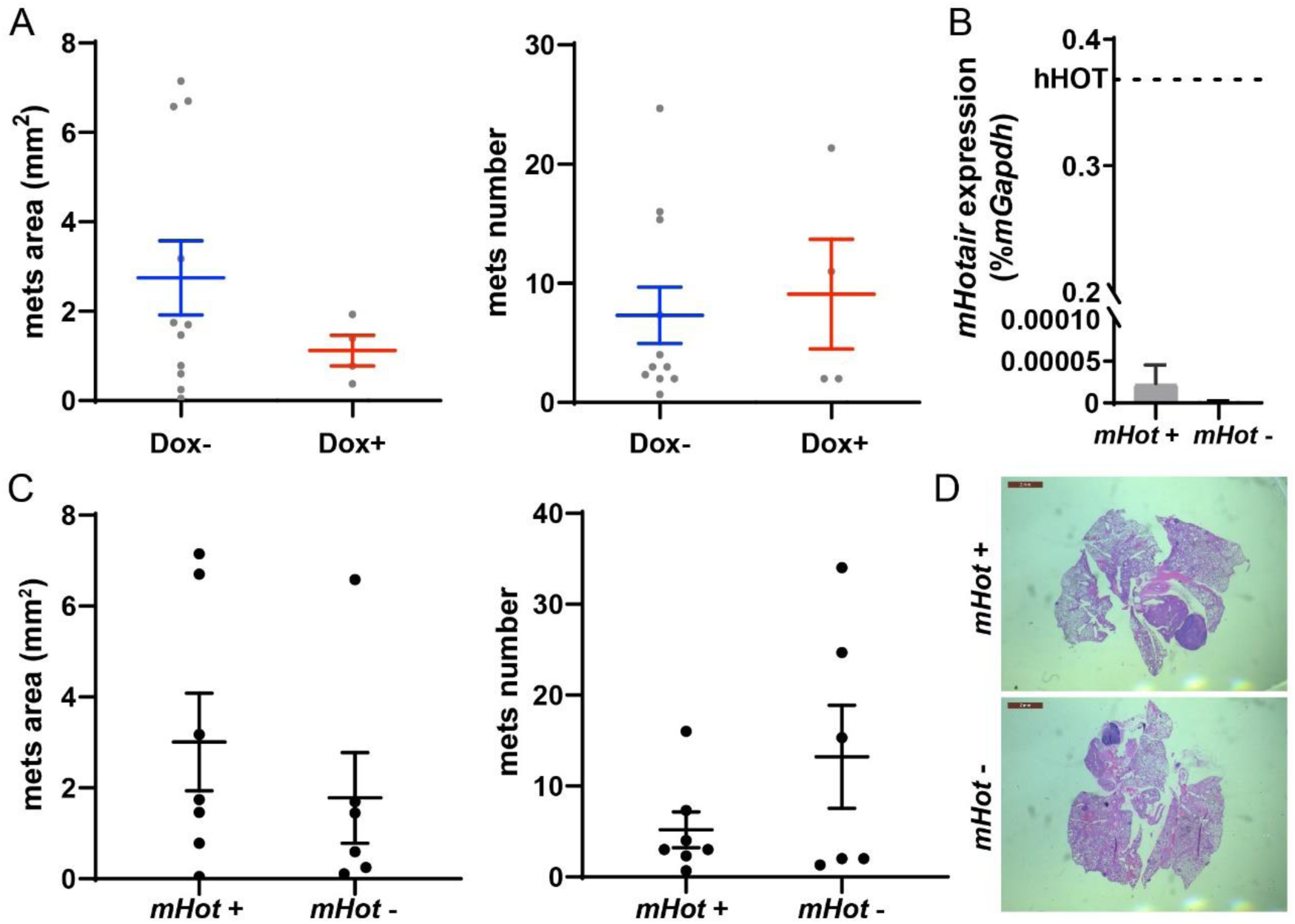
Dox itself or endogenous mouse *Hotair*, which expresses at very low level, have no significant effect in breast tumor metastasis. A: Dox has no significant influence in tumor progression of MMTV-PyMT mice. MMTV-PyMT mice were treated with or without Dox for about 100-120 days. Quantification of lung metastases was based on hematoxylin and eosin (H&E) staining. Metastases area and numbers in sections were calculated by ImageJ. Dashes represent means ± SEM, n = 11&4. Spots represent every single value. Student t-test was performed and showed no significant difference between Dox-and Dox+ group. B: Endogenous mouse *Hotair* expression is very low compared to transformed human HOTAIR. qRT-PCR was performed to measure endogenous mouse *Hotair* expression in *mHotair*+ and *mHotair*- MMTV-PyMT mice. Dashed line demonstrated human HOTAIR level in iHOT-PyMT mice without Dox treated (same as Fig 2B). Values are means ± SEM, n = 3&2. mHot is short for mouse Hotair, and hHOT is short for human HOTAIR. C-D: Endogenous mouse *Hotair* knockout has no significant influence in breast tumor metastasis in MMTV-PyMT mice. Quantification of lung metastases was based on H&E staining. Metastases area and numbers in sections were calculated by ImageJ. Dashes represent means ± SEM, n = 6&7. Spots represent every single value. Student t-test was performed and showed no significant difference between *mHotair*+ and *mHotair*-group. Scale bar = 2 mm.

**Supp 3:**
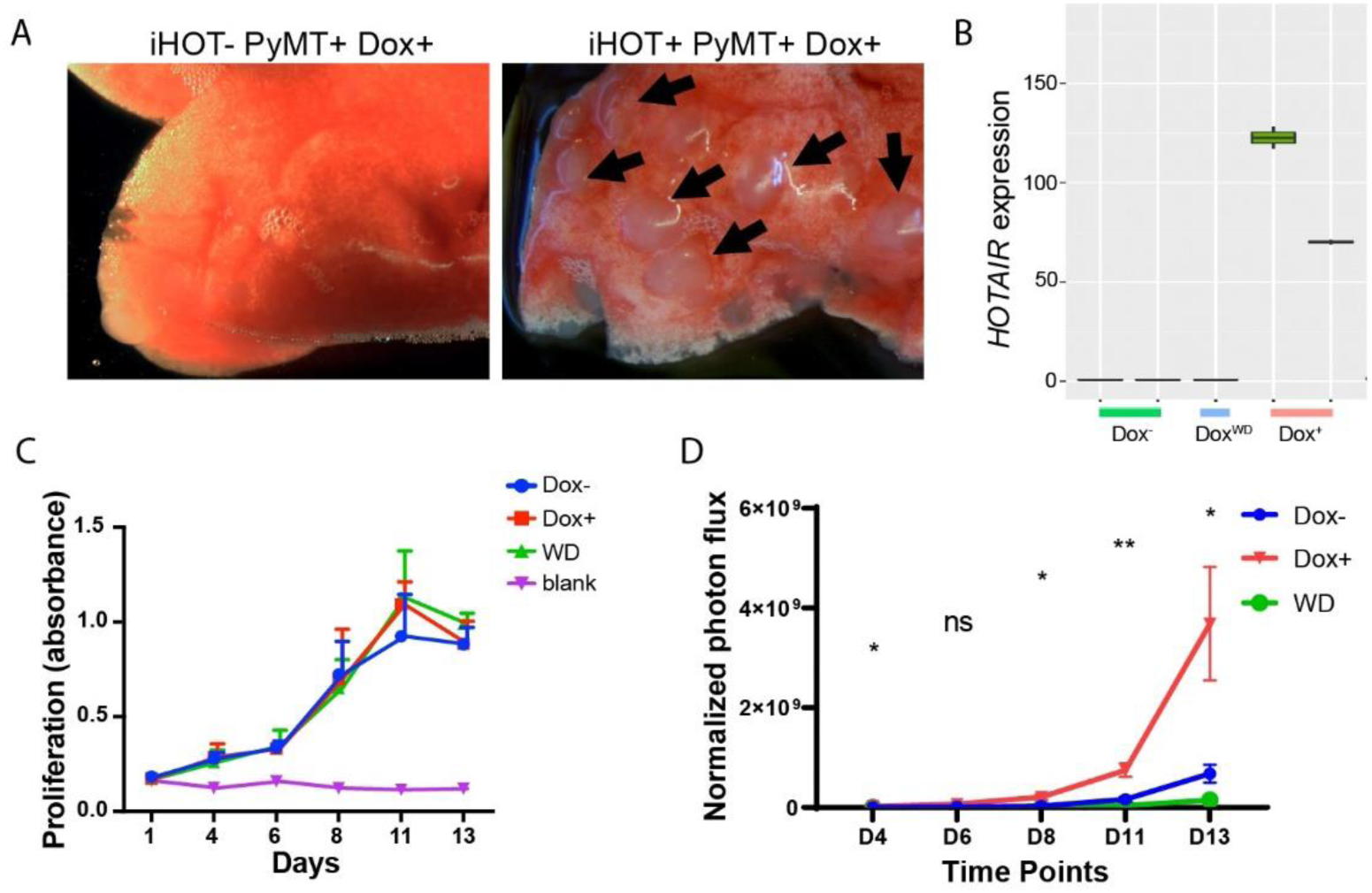
Induced HOTAIR promotes tumour cell metastasis but have no significant effect on cell proliferation. A. Images of HOTAIR overexpressing iHOT-PyMT mouse lungs under a dissecting microscope. Arrows indicate metastatic tumors colonizing lung tissue. B. Expression of HOTAIR level (FPKM value) in RNA-seq data. C. Dox^-^, Dox^+^, Dox withdraw treatment does not have a statistically significant effect on cellular proliferation. Blank controls are cells without substrate. D: Another batch of tail vein injection assay. Dox treated iHOT^+^ cells overexpressing HOTAIR display higher rates of lung colonization compare to the Dox^-^ group (n = 5-6). Student t-test was performed between Dox+ and Dox-group at each time point, *P<0.05, **P<0.05.

**Supp 4:**
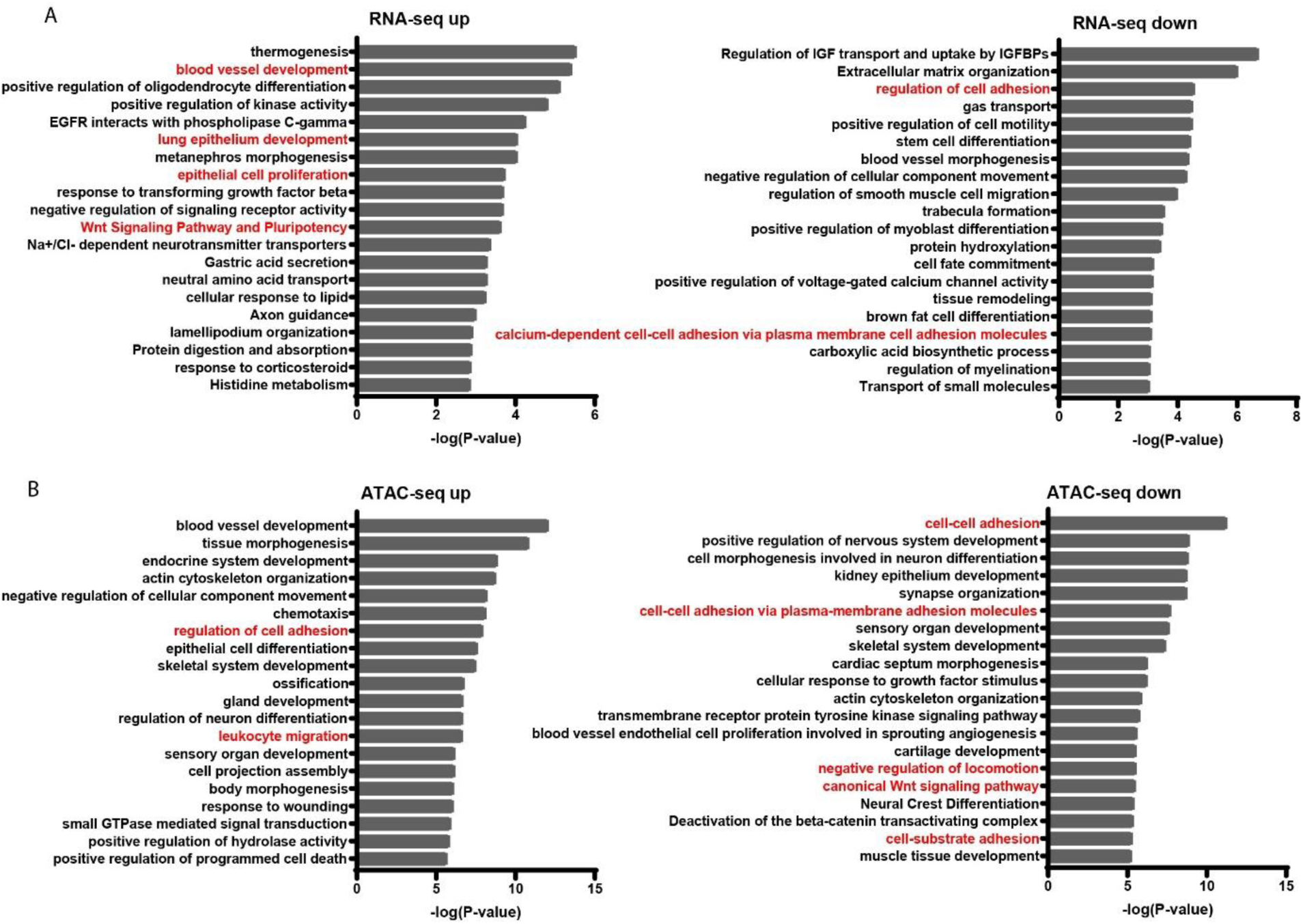
GO enrichment analysis of significant differential analysis for RNA-seq and ATAC-seq data. A. GO enrichment analysis of differentially expressed genes by RNA-Seq in Dox^-^ and Dox^+^ iHOT cells. B. GO enrichment analysis of differential peaks associated genes by ATAC-Seq in Dox^-^ and Dox^+^ iHOT cells. Terms highlighted in red are examples of terms related to cancer cell metastasis.

